# Effect of spatial overdispersion on confidence intervals for population density estimated by spatial capture–recapture

**DOI:** 10.1101/2024.03.12.584742

**Authors:** Murray G. Efford, David Fletcher

**Author notes:** Corresponding author: Murray Efford.

## Abstract

Spatially explicit capture–recapture models are used widely to estimate the density of animal populations. The population is represented by an inhomogeneous Poisson point process, where each point is the activity centre of an individual and density corresponds to the intensity surface. Estimates of density that assume a homogeneous model (‘average density’) are robust to unmodelled inhomogeneity, and the coverage of confidence intervals is good when the intensity surface does not change, even if it is quite uneven. However, coverage is poor when the intensity surface differs among realisations. Practical examples include populations with dynamic social aggregation, and the population in a region sampled using small detector arrays. Poor coverage results from overdispersion of the number of detected individuals; the number is Poisson when the intensity surface is static, but stochasticity leads to extra-Poisson variation.

We investigated overdispersion from three point processes with a stochastic intensity surface (Thomas cluster process, random habitat mosaic and log-Gaussian Cox process). A previously proposed correction for overdispersion performed poorly. The problem is lessened by assuming population size to be fixed, but this assumption cannot be justified for common study designs. Rigorous correction for spatial overdispersion requires either prior knowledge of the generating process or replicated and representative sampling. When the generating process is known, variation in a new scalar measure of local density predicts overdispersion. Otherwise, overdispersion may be estimated empirically from the numbers detected on independent detector arrays.

## Introduction

Spatially explicit capture–recapture (SECR) is a method for estimating the density of an animal population by modelling observations of marked individuals at detectors in known locations. Detectors may be traps, automatic cameras, DNA hair snares or other devices. Individual marks may be natural (e.g., microsatellite DNA or pelage patterns) or applied on first capture (e.g., numbered tags or bands). The state model is the spatial distribution of animal activity centres (AC). Activity centres are not observed directly, but are modelled as a realisation of a spatial point process. Capture–recapture sampling (the observation process) provides spatial data on a sample of animals. Parameters of the state model (specifically, the intensity surface of the point process) are estimated jointly with parameters of the spatial observation process by maximizing the marginal likelihood (Borchers & Efford, 2008) or in a Bayesian framework by data augmentation (Royle et al., 2014) or using the semi-complete data likelihood (Zhang et al., 2023).

The inhomogeneous Poisson process (IHPP) serves as a general model for the distribution of AC in SECR (Borchers & Efford, 2008). Animal population density is represented by the intensity surface. Spatial variation in density may be explained in part by covariates such as habitat type or proximity to roads or other hazards. However, the cause of much variation is unknown and possibly unknowable: the distribution of AC may bear the imprint of stochastic demography, social aggregation, transient resources, harvesting, local catastrophes or colonisation processes. It is natural then to report the ‘average’ density *D*^^^ estimated by fitting a uniform-density (homogeneous) model, even if the true density is likely to be inhomogeneous.

We consider the consequences of misspecifying the density model as homogeneous when the true intensity is inhomogeneous and possibly dynamic. The primary concern, that average density estimates will be biased by inhomogeneity, turns out to be unwarranted when detection does not vary over space. A secondary concern is that unmodelled inhomogeneity of density could cause underestimation of sampling variance due to overdispersion in the number of detected individuals *n*. Confidence intervals (CI) for density would then be misleadingly narrow and inference unreliable. Whether overdispersion does arise and cause these effects depends on the scope of inference and the underlying biology, as we hope to make clear.

Our starting point is a static IHPP in which both the intensity surface and the locations of detectors are fixed. Static inhomogeneity does not result in overdispersion in *n* regardless of the unevenness of the surface, and CI that assume Poisson *n* are reliable. We justify this assertion in a later section.

Bischof et al. (2020) drew attention to poor coverage of confidence intervals for SECR estimates of average density when animals are socially aggregated in family groups or packs. Coverage improved when they applied a variance inflation factor based on the number of individuals per detector. Social aggregation implies spatial clustering of AC, and a snapshot of a clustered distribution is not distinguishable from one that is a realisation of a static IHPP with local ‘peaks’ of intensity (e.g., Diggle et al., 2013). Why then should this scenario result in overdispersion and poor CI coverage? The key is that Bischof et al. (2020) related their simulated estimates not to the local intensity surface of a particular static IHPP near their detectors, but to a global density of which that was one realisation. Clustering due to animal behaviour is intuitively dynamic: given time, the same individuals may form different groupings in different locations. A plausible reference distribution thus includes redistribution across the landscape rather than just a snapshot.

This leads us to the concept of a stochastic-intensity IHPP, otherwise known as a Cox process (e.g., Møller & Waagepetersen, 2004; Chiu et al., 2013). The intensity surface of a Cox process varies among realisations. In a widely used clustering model, the Thomas process, clusters are seeded by randomly locating ‘parent’ points in the plane and each parent gives rise to a Poisson number of ‘offspring’ scattered about the parent according to a circular bivariate normal distribution (Fig. 1 and Appendix S1). Each realised distribution of parents corresponds to a static IHPP intensity surface (Møller & Waagepetersen, 2004 p. 61), but clusters are spatially unmoored and not tied to persistent landscape features. Cox processes arise mathematically in many other ways. AC may be restricted to one phase of a binary habitat mosaic whose configuration differs between realisations; we refer to this as a ‘random habitat’ (RH) process. The log-Gaussian Cox process (LGCP) is a popular point process model (Møller et al., 1998; Diggle et al., 2013) that was used by Dupont et al. (2021) to simulate AC for SECR.

**Figure 1:**
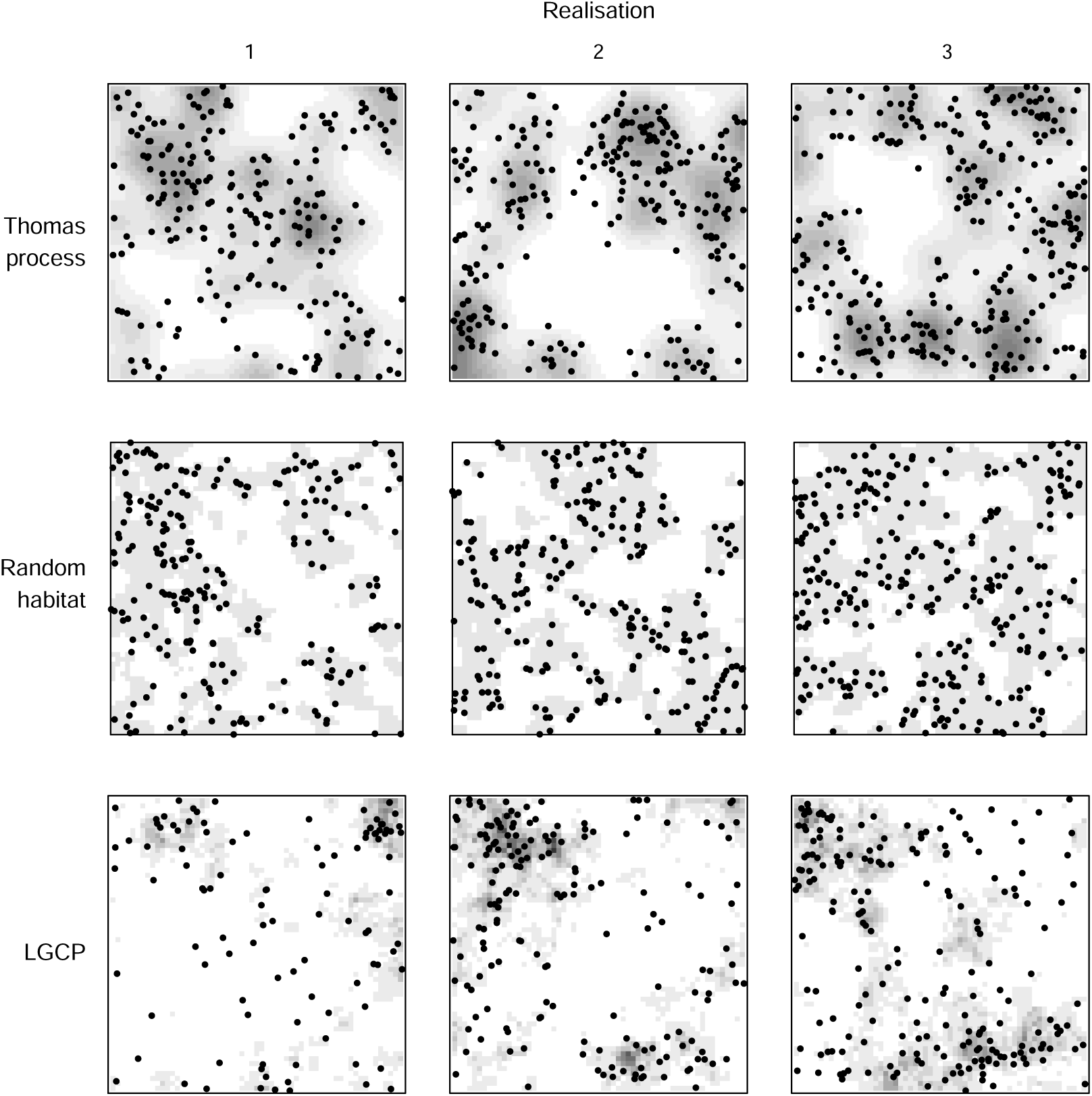
Distributions of activity centres generated by various Cox processes (stochastic-intensity IHPP) with expected number 256 in a square 30σ × 30σ (σ defined in text). Columns indicate different realisations with the same parameter values. Grey shading indicates intensity of IHPP. See Appendix S1 for more. (a) Thomas process clustering: each uniformly distributed ‘parent’ (not shown) gives rise to a Poisson number of offspring (expectation μ = 8) that follow circular bivariate normal distribution about the parent (scale 2σ). (b) Random habitat mosaic (fragmentation parameter *p* = 0.5, fraction *f* = 0.5). (c) Log-Gaussian Cox Process with variance *V* = 1.0 on log scale; exponential spatial covariance scale = 5σ.

Reliable estimates of the sampling variance, and therefore of confidence intervals for *D*^^^ relative to the global density, might in principle be obtained by expanding the fitted model to include the generating Cox process for AC. That approach was applied in the context of distance sampling by Waagepetersen & Schweder (2006). It is unlikely to be practical in SECR: the generating process is usually unknown, and we would stand little chance of fitting it with data from a single realisation even if the process were known.

A model-free approach to overdispersion has entered non-spatial capture–recapture studies (Burnham et al., 1987, pp. 243–246; Lebreton et al., 1992; Anderson et al., 1994; White & Cooch, 2017) from generalized linear modelling (Wedderburn, 1974). An empirical estimate of the overdispersion ratio *c*, based on a goodness-of-fit statistic, may be used to adjust variances and achieve near-nominal coverage of confidence intervals. Thus for a parameter ψ, var_adj_(ψ^^^) = *c*^ var(ψ^^^), usually before back-transformation from a ‘link’ scale such as logit or log.

The focus for non-spatial capture–recapture is on non-independence and heterogeneity among animals, for which the observed and expected frequencies of capture histories provide the required empirical index *c*^. For spatial overdispersion we require a specifically spatial measure.

In this paper we describe properties of the static and stochastic IHPP models for AC and use simulation to demonstrate the effects of static and dynamic inhomogeneity on density estimates. We also consider the practical challenge of adjusting for overdispersion in *n*. We simulate the variance inflation factor of Bischof et al. (2020) for a range of Cox processes and find that it fails to correct for overdispersion in *n*. Empirical estimation of overdispersion is possible with replicated samples from the Cox process. Spatially replicated small arrays of detectors (‘subgrids’) have been used in SECR to estimate density in a larger region (e.g., Humm et al., 2017; Clark, 2019; Howe et al., 2022). This is akin to distance sampling, in which each line or point defines a plot (Buckland et al. 2001). We can view the density at each subgrid as resulting from one realisation of a Cox process whose global mean is the required regional density. Given some design assumptions (independence and representativeness of subgrids) the overdispersion of the combined *n* may be estimated empirically from variation in its subgrid-specific components, and used to obtain confidence intervals with near-nominal coverage. We demonstrate this by simulation for a simple scenario and reconcile our results with those of Howe et al. (2022).

## Background

For specificity we focus on closed SECR models and observations at point detectors such as automatic cameras. We use ***ω*** for the set of *n* observed spatial detection histories, and ø and θ for the parameter vectors of the state and observation models respectively. The likelihood has the general form *L*(ø, θ|***ω***) = Pr(*n*|ø, θ) Pr(***ω***|*n*, ϕ, θ) (Borchers & Efford, 2008). The intensity surface of the IHPP for AC is a function of location, *D*(**x**; ø), where **x** represents the Cartesian coordinates *x, y*. We denote the estimate of density from a homogeneous model by *D*^^^ (without dependence on **x**). Density may be estimated from a homogeneous model in two stages, first estimating θ by maximizing the second factor in the likelihood conditional on *n*, and then computing the Horvitz-Thompson-like estimator *D*^^^ = *n*/*a*(θ^^^) (Borchers & Efford, 2008). The effective sampling area *a* is a function of the detection parameters *a*(θ) = *p*_·_ (**x**; θ) *d***x**, where *p*_·_ (**x**; θ) models the overall probability of detection for an AC at **x** and integration is over all habitat in the plane.

Spatial capture–recapture models use a distance-dependent detection function that takes many possible forms. Let *d*_*k*_ = |**x** − **x**_*k*_ | be the distance between an individual’s AC (**x**) and the location of detector *k* (**x**_*k*_). We used a hazard half-normal function in which the modelled hazard of detection in detector *k* is λ_*k*_ = λ_0_ exp[−*d*^2^/(2σ^2^)]. The overall probability an individual with AC at **x** is detected at least once at one of *K* detectors operated for *S* occasions is then *p*_·_ (**x**; θ) = 1 − exp[−*S* _*K*_ λ_*k*_]. This function defines a surface that drops to zero away from the detectors.

As in distance sampling (e.g., Buckland et al., 2001), the variance of *D*^^^ may be approximated by the delta method as

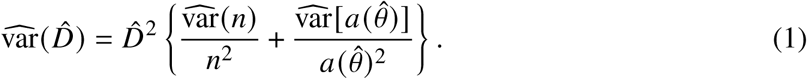

Maximization of the conditional likelihood provides an estimate of θ and hence var[*a*(θ^^^)]. This term is often smaller than the first. If *n* is Poisson then the first term inside the braces simplifies to 1/*n*. However, to assume wrongly that *n* is Poisson risks biasing estimates of var(*D*^^^).

### Static IHPP

For an intensity surface that is flat or a deterministic function of spatial covariates or trend, the expected number of AC in a region *A* is E[*N* (*A*)] = _*A*_ *D*(**x**; ø) *d***x**, and from the properties of an IHPP, *N* (*A*) is Poisson with variance E[*N* (*A*)] (e.g., Diggle, 2003 p. 67). The spatial detection process applies location-dependent thinning to the distribution of activity centres. Thinning is independent for the detector types we consider, but not for single-catch traps in which a maximum of one individual may be caught on any occasion. The expected number of animals detected *n* is an integral over space of the product of density and a location-dependent detection function, i.e.

The probability of detection is assumed to approach zero at large distances from the detectors. Consider an area *A*_*w*_ that includes the detectors and all habitat within a buffer of width *w*. Then E(*n*) approaches a constant as *w* increases, and beyond some *w*^′^ further change in E(*n*) is negligible. This is convenient for computation because numerical integration may be restricted to any arbitrary area *A*_*w*_ where *w* > *w*^′^, without causing bias in density estimates.

If AC are distributed as an inhomogeneous Poisson process and detections are independent, then the locations of detected AC are also inhomogeneous Poisson, by the properties of independently thinned point processes (Chiu et al., 2013). It follows that the number of animals detected is Poisson, with variance var(*n*) = E(*n*) (Borchers & Efford, 2008). Thus inhomogeneity of density does not in itself imply overdispersion.

The population size *N* (*A*) in an area *A* does not appear in the likelihood for the IHPP SECR model. However, in many implementations *N* (*A*) is considered fixed rather than random and does appear in the likelihood, which then depends on *A*. We doubt the general applicability of such models, as discussed later, but provide results here that complement those for Poisson *N* (*A*). The distribution of AC with fixed *N* (*A*) is a binomial point process (Illian et al., 2008) conditional on *N* (*A*). The expected value of *n* is unchanged from its Poisson expectation, but the variance of *n* is reduced. Specifically, var(*n*) = E(*n*)(1 − *p*) where *p* = _*A*_ *p*_·_ (**x**; θ)*D*(**x**; ø) *d***x**/ _*A*_ *D*(**x**; ø) *d***x** (Illian et al., 2008, p. 107). If density is homogeneous this simplifies to var(*n*) = E(*n*)(1 − *a*(θ)/|*A*|) where |*A*| is the area of *A*.

### Stochastic IHPP

The intensity of a Cox process (an IHPP with stochastic intensity) is a spatial random variable. From here on we use *D*(**x**) to mean this variable, and omit the parameter vector to lighten the notation. The specification of *D*(**x**) as a Cox process is flexible and requires parameters additional to ø that we consider only in passing because, while they are used in simulation, they cannot easily be estimated from SECR data.

We considered three Cox processes:

a. the Thomas cluster process, a variety of Neyman-Scott process, for which the initial distribution of parents determines the intensity surface (see Introduction),
b. a random habitat mosaic (RH) in which habitat pixels with fixed positive density are clustered in quasi-realistic random configurations within a known fraction of the landscape and intervening areas have zero density, and
c. the Log-Gaussian Cox process (LGCP), in which the intensity is a continuously varying Gaussian random field.

Each Cox process is realised in two stages: first a random intensity surface is generated and then AC locations are generated from the intensity surface as an inhomogeneous Poisson point pattern. Examples are shown in Fig. 1.

### Global and local density

We next consider how to describe the intensity surface of a Cox process for AC in a way that is relevant to SECR. The global ‘average’ density is simply the expected value of *D*(**x**). Stochastic intensity surfaces may include a deterministic trend (e.g., Johnson et al., 2010), but for simplicity we assume each process is stationary. In particular, we assume that the expected-density surface is flat, so that we can use the simpler notation for global density μ_*D*_ ≡ E[*D*(**x**)].

SECR samples AC from a local ‘window’ of the global surface. The window is defined probabilistically by the overall-detection function *p*_·_ (**x**; θ). We therefore define the realised local density in the vicinity of detectors *D*_*a*_ to be an integral of the density surface weighted by the probability of detection:

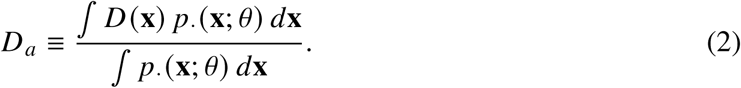

This is a scalar random variable. *D*_*a*_ for known θ may be computed directly from a particular intensity surface *D*(**x**). Local density differs among realisations of the intensity surface. It can be shown that E(*D*_*a*_) = μ_*D*_, given that μ_*D*_ is independent of location. The magnitude of var(*D*_*a*_) depends on both the Cox process and the window function *p*_·_ (**x**; θ), which in turn depends on the detector array, the duration of sampling, and θ.

### Spatial overdispersion

Underestimation of the global sampling variance of *D*^^^ using Equation 1 results directly from excess variation in *n* relative to the fitted model. Confidence intervals based on such estimates will therefore have poor coverage of the global density, as we demonstrate later by simulation.

Coverage can be restored with a variance inflation factor incorporating the overdispersion in *n*. We define the amount of overdispersion in *n* by the ratio *c*_*n*_ = var(*n*)/var_0_(*n*) using var_0_(*n*) to denote the variance of *n* under the fitted model. We assume a Poisson model for *n* and hence var_0_(*n*) = E(*n*), except when addressing fixed population size *N* (*A*) for which *n* is binomial.

We show in Appendix S2 that var(*n*) for a stochastic-intensity IHPP may be written as var(*n*) = E(*n*) + *a*(θ)^2^ var(*D*_*a*_) where E(*n*) = *a*(θ)E(*D*_*a*_), and hence

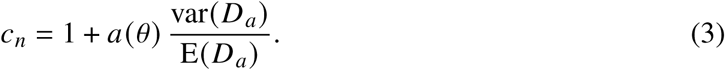

The second term indicates the overdispersion in *n* that is caused by stochasticity in the intensity of the IHPP.

Equation (3) allows us to predict *c*_*n*_ from θ and moments of the distribution of *D*_*a*_. Simulation is at present the only practical and general method for computing these moments (mean and variance), but we need to simulate only the intensity-generating phase of a Cox process, i.e. the distribution of parents for the Thomas process, habitat mosaics for RH, and random fields for LGCP. Estimates of *c*_*n*_ based on var(*D*_*a*_) and known θ were indistinguishable in practice from those calculated by simulating the full sampling process to obtain var(*n*) (Appendix S3).

### Correction for overdispersion

Johnson et al. (2010) constructed a measure of spatial overdispersion from the observed and expected numbers of detections on distance-sampling transects, but coverage of adjusted confidence intervals for density remained poor in some scenarios. Bischof et al. (2020) formed a similar measure from the observed (*n*_*k*_) and expected (E(*n*_*k*_)) numbers of SECR detections per detector *k*:

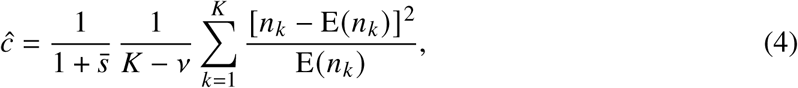

where *K* is the number of detectors, *v* is the number of estimated parameters, and *s^-^* = 1/K Σ^K^_K=1_ [*n*_*k*_ − E(*n*_*k*_)]/E(*n*_*k*_) is a correction for sparse data due to Fletcher (2012). The estimator assumes each *n*_*k*_ is drawn from a detector-specific Poisson distribution. Bischof et al. (2020) applied *c*^ as a variance inflation factor to simulated data from clustered AC, with mixed results.

Overdispersion may be estimated for *J* independent spatial samples from a stochastic IHPP from the same formula as Equation 4, but with the per-detector counts *n*_*k*_ replaced by per-array counts *n*_*j*_. We label this estimator *c*^_*n*_. For sampling with independent equal-sized subgrids operated for the same duration and a homogeneous density model, E(*n*_*j*_) = *n*/*J* and *s^-^* = 0. Then *c*^_*n*_ is simply the ratio of the sample variance of the *n* _*j*_ to their mean:

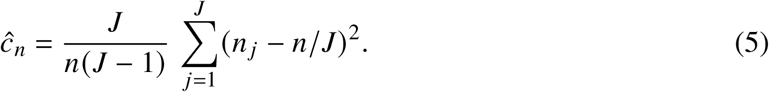

We have not explored possible weighting for variation among arrays (cf Fewster et al. 2009).

## Methods

We used simulation to evaluate the performance of a homogeneous-density model fitted to SECR data in which AC followed the three Cox processes each with varying levels of stochasticity. The sampling design consisted of a 12 × 12 array of binary proximity detectors (*sensu* Efford et al., 2009) operated for 5 occasions; detector spacing was 2σ where σ was the spatial scale parameter of a hazard halfnormal detection function and λ_0_ = 0.5. AC (expected number 256) were distributed as detailed below within an area extending *w* = 4σ beyond the detectors. SECR detection histories were generated with the function ‘sim.capthist’ of R package ‘secr’. Each scenario was replicated 1000 times. We repeated each of the main simulations with a conditional form of the Cox process i.e. using fixed *N* (*A*) for both the generating process and the fitted model.

### Cox processes for AC

#### Thomas cluster process

Cluster process simulations used the ‘spatstat’ function rThomas (Baddeley et al. 2015), parameterized by the overall density, the mean number per cluster μ ∈ {1, 2, 4, 8, 16, 32} and the scale of spatial spread within clusters (2σ where σ was the scale of the halfnormal SECR detection model). Bischof et al. (2020) simulated a fixed number of clusters and fixed number in each cluster, with zero within-cluster dispersion. We were able to simulate more realistic clustering (varying number and dispersion).

#### Random habitat

Random habitat patches comprising a varying fraction *f* of *A*_*w*_ were generated by the modified random cluster algorithm of Saura and Martínez-Millán (2000). The patch-fragmentation parameter (‘initial probability’) was set to 0.5. The expected density is E[*D*(**x**)] = *f D*_*p*_ where *D*_*p*_ is the density within habitat patches; *D*_*p*_ was therefore adjusted upwards by 1/ *f* to maintain constant overall density.

#### Log-Gaussian Cox process

For LGCP the log-intensity surface was modelled as a Gaussian random field; the field is parameterized by its mean, variance and scale of spatial covariance. For overall point density *D* and Gaussian variance *V*, the required mean is μ = log(*D*) − *V*/2. We used the function rLGCP in the package ‘spatstat’ (Baddeley et al., 2015). The spatial covariance function was exponential.

Simulations used six levels of variance (*V* ∈ {0, 0.125, 0.25, 0.5, 0.75, 1.0}) and spatial scale 10σ.

### Parameter estimation

A homogeneous-density SECR model was fitted to each dataset by maximizing the full likelihood (Borchers & Efford, 2008). Area *A*_*w*_ extending 4σ beyond the detectors was discretized as cells of width 0.5σ. E(*n*) ≈ 180.7 and *p* ≡ *a*/|*A*_*w*_ | ≈ 0.7058 for all scenarios, where |*A*_*w*_ | is the area of *A*_*w*_ and *p* is the probability an AC in *A*_*w*_ is detected. Relative bias and confidence interval coverage were computed with respect to both local and global definitions of true density, both with and without adjustment for overdispersion when *c*^ > 1.

Computations used R packages secr 5.2.2 (Efford, 2025a), secrdesign 2.9.3 (Efford, 2025b) and spatstat 3.3-3 (Baddeley et al., 2015). Documented code and results are included in an *ad hoc* R package ‘overdispsim’ that is available on GitHub (Efford, 2025c).

### Empirical overdispersion

To evaluate *c*^ as an estimate of *c*_*n*_ we conducted further simulations of AC distributed according to the three Cox processes, followed by spatial sampling to obtain *n*, but without the time-consuming step of fitting an SECR model. These simulations were faster because model fitting was not required and the number of replicates was therefore increased to 10000.

### Spatial replication

We performed limited simulations to contrast the effects of static and stochastic IHPP on SECR estimates of regional density from replicated subgrids. Howe et al. (2022) simulated the use of replicated arrays of DNA hair snags to sample populations of black bear (*Ursus americanus*). Our simulations followed their Scenario 2 to allow results to be compared directly. Two 5 × 8 subgrids (2-km detector spacing) were notionally placed in each of three density zones (6, 12 or 18 bears per 100 km2) extending 15 km from the detectors. Following Howe et al. (2022) we used the detection function *g*(*d*_*k*_) = *g*_0_ exp[−*d*^2^/(2σ^2^)] where *g* refers to probability of detection rather than hazard; parameters were constant (*g*_0_ = 0.3, σ = 1.5 km, 6 sampling occasions). We fitted a homogeneous model to the pooled data to estimate regional density. Overdispersion was estimated from the numbers of individuals per subgrid as described in ‘Correction for overdispersion’. We evaluated relative bias and the coverage of 95% confidence intervals for regional density, with and without correction for overdispersion, over 1000 replicates. AC patterns under the original scenario were realisations of a static IHPP. For comparison we also simulated AC from a stochastic IHPP by randomly selecting the density at each subgrid from the three possibilities (6, 12 or 18 bears per 100 km2), with replacement. Simulations were also repeated with a larger sample of subgrids (18).

## Results

Simulations confirmed the lack of bias in estimates of both local and global density from a homogeneous model even when the generating process for AC was highly clumped and stochastic (Fig. 2, detailed results in Appendix S4).

**Figure 2:**
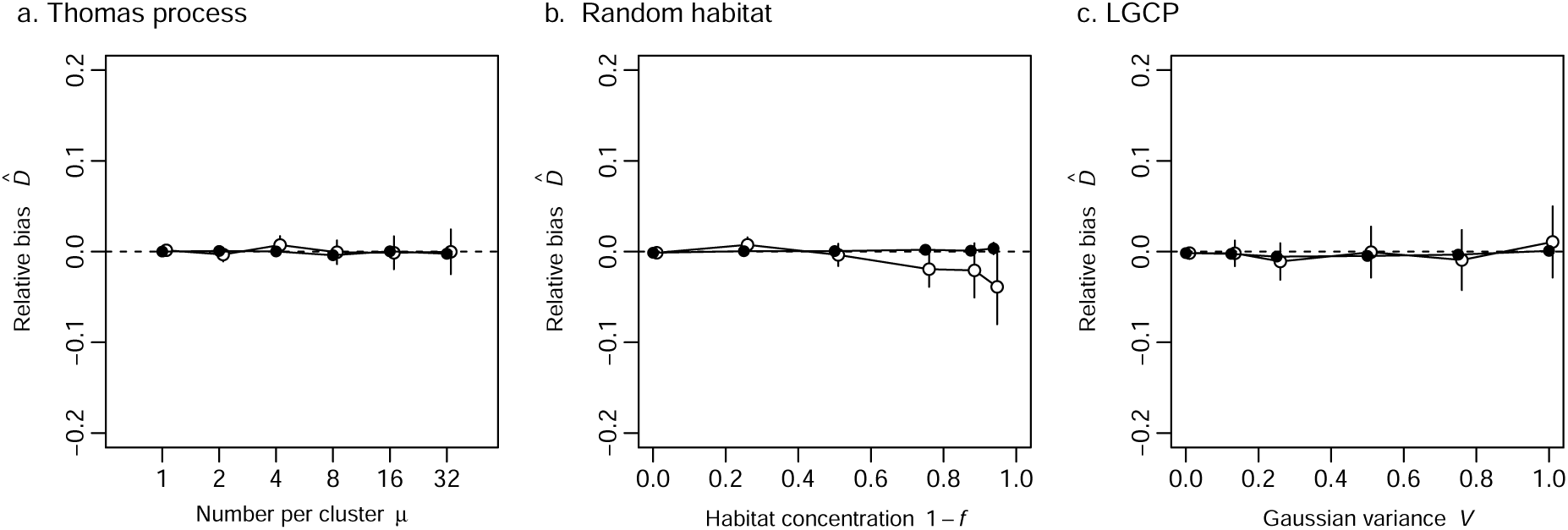
Relative bias of density (*D*^^^) estimated by fitting a homogeneous model when AC followed one of three Cox processes. (a) Thomas cluster process with varying expected number per cluster μ, (b) random habitat patches occupying fraction *f* of landscape, and (c) log-Gaussian Cox process with variance *V*. Estimates were related to either the global density (◦) or the detection-weighted local density (•) (see text). 1000 simulations per scenario; 95% confidence intervals.

For the same scenarios we next compared the empirical (simulation-based) estimate of overdispersion *c*_*n*_ to the variance inflation factor *c*^ computed by Bischof et al. (2020). Although both increased with the underlying variance, the overdispersion due to each Cox process greatly exceeded *c*^ (Fig. 3). See Appendix S3 for results from a wider range of scenarios.

**Figure 3:**
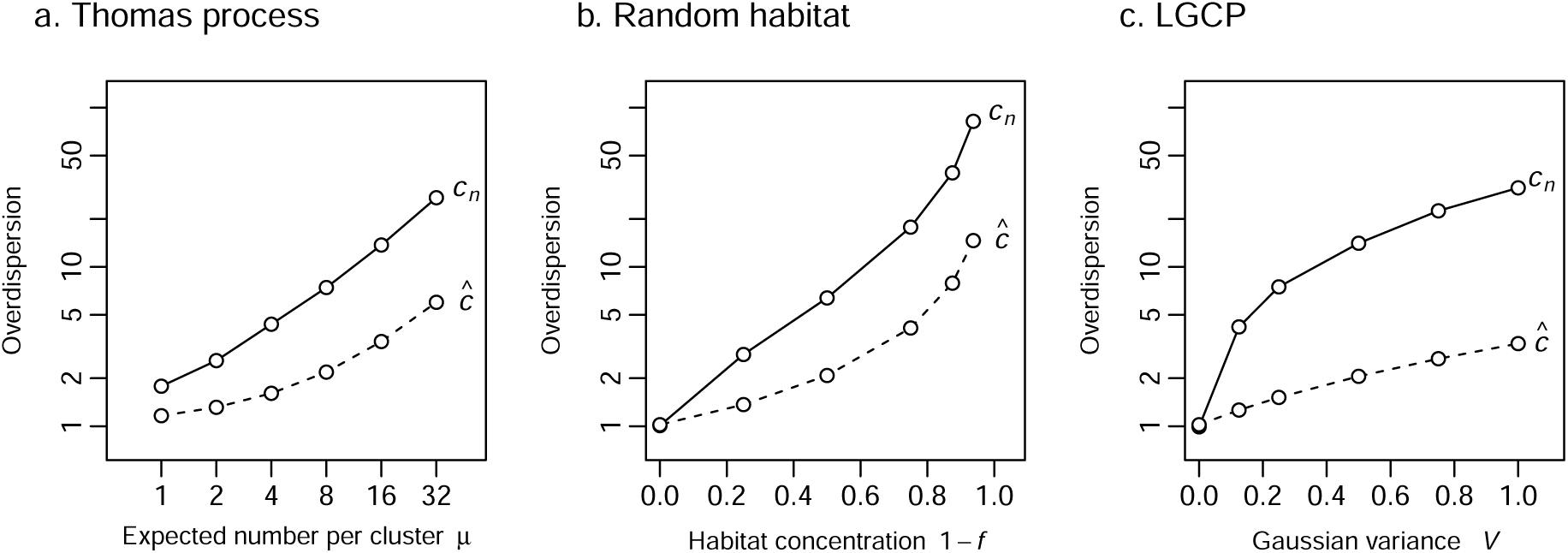
Simulated overdispersion of number detected *c*_*n*_ and the variance inflation factor of Bischof et al. (2020) for three Cox processes. (a) Thomas cluster process with varying expected number per cluster μ, (b) random habitat patches occupying fraction *f* of landscape, and (c) log-Gaussian Cox process with variance *V*. 10000 simulations per scenario.

Confidence intervals for *D*^^^ achieved the nominal coverage of local density, but failed to cover the global density of AC from Cox or cluster processes (Fig. 4 and Appendix S4). The term for var(*n*) in Equation (1) contributed > 98% of var(*D*) across all simulation scenarios.

**Figure 4:**
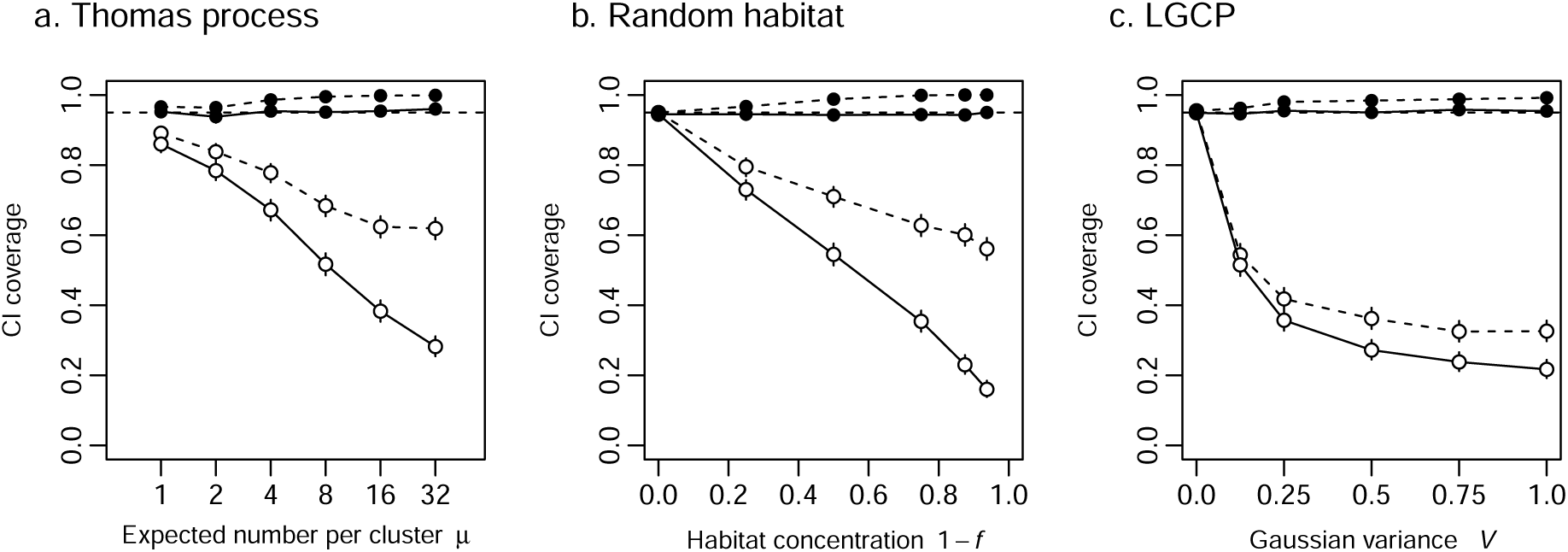
Coverage of 95% confidence intervals for *D*^^^ when AC followed one of three Cox processes. (a) Thomas cluster process with varying expected number per cluster μ, (b) random habitat patches occupying fraction *f* of landscape, and (c) log-Gaussian Cox process with variance *V*. Estimates were related to either the global density (◦) or the detection-weighted local density (•). Intervals were unadjusted (solid line) or adjusted with the variance inflation factor of Bischof et al. (2020) (dashed line). 1000 simulations per scenario; 95% confidence intervals.

When we applied the correction of Bischof et al. (2020) the results were not encouraging. Interval coverage invariably increased (Fig. 4), but the adjustment fell well short of providing nominal coverage for all stochastic scenarios. Inflating the variance resulted in intervals for local density that were unnecessarily wide, with coverage approaching 100% (Fig. 4).

Simulation results are shown in Fig. 5 for the fixed-*N* (*A*) (i.e. conditional) equivalents of Fig. 4.

**Figure 5:**
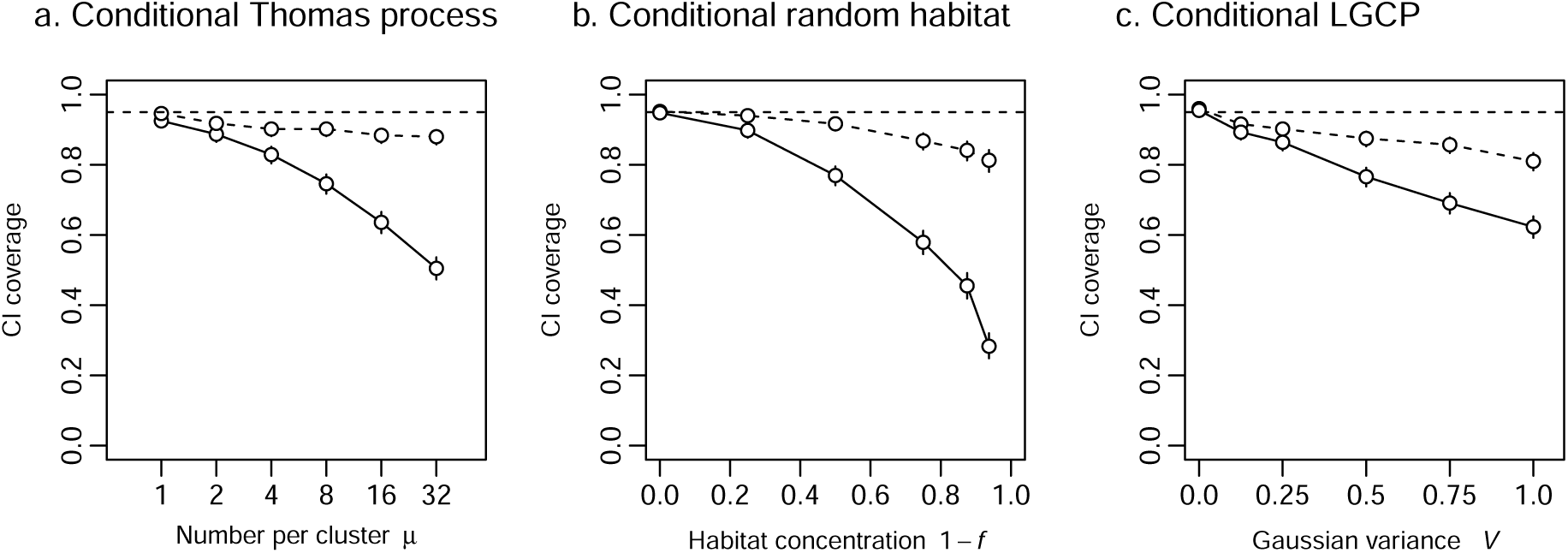
Coverage of 95% confidence intervals for *D*^^^ as function of clumping when *N* (*A*) was fixed and fitted *n* binomial. Intervals were unadjusted (solid line) or adjusted with the variance inflation factor of Bischof et al. (2020) (dashed line). (a) Thomas cluster process with varying expected number per cluster μ, (b) random habitat patches occupying fraction *f* of landscape, and (c) log-Gaussian Cox process with variance *V*. 1000 simulations per scenario; 95% confidence intervals.

### Spatial replication

Simulations of sampling with multiple subgrids showed good coverage of confidence intervals for regional density when the intensity was static relative to the detectors, but coverage was poor when the intensity was stochastic (Table 1). Adjustment in the ‘static’ case was unwarranted and resulted in excessive coverage approaching 100%. When the intensity was stochastic, inflating variances by the factor *c*^_*n*_ gave CI coverage that was closer to nominal, especially in the scenario with more subgrids (Table 1).

**Table 1:** Simulated SECR estimates of regional population density using *J* replicate subgrids when the intensity of the inhomogeneous Poisson process is static or stochastic. Density estimates are summarised as relative bias (RB), relative standard error (RSE) and the coverage of 95% confidence intervals (COV). The estimate of overdispersion *c*^_*n*_ is a variance inflation factor used to compute RSE_adj_, and coverage COV_adj_; adjustment is inappropriate when the intensity is static (*). SE in parentheses. All SE(RSE) < 0.0005 and SE(RSEadj) < 0.002. Average of 1000 replicates.

| IHPP | $J$ | RB | RSE | COV | $\hat{c}_n$ | $\text{RSE}_{\text{adj}}$ | $\text{COV}_{\text{adj}}$ |
| --- | --- | --- | --- | --- | --- | --- | --- |
| static | 6 | -0.003 (0.002) | 0.076 | 0.961 | 7.06 (0.07) | 0.202* | 1.000* |
|  | 18 | -0.001 (0.001) | 0.044 | 0.945 | 6.35 (0.03) | 0.111* | 1.000* |
| stochastic | 6 | -0.002 (0.006) | 0.077 | 0.589 | 6.16 (0.10) | 0.188 | 0.894 |
|  | 18 | 0.002 (0.003) | 0.044 | 0.566 | 6.11 (0.05) | 0.108 | 0.932 |

### Overdispersion in detection

Our focus has been on the AC point process and its effect on the overdispersion of *n* and the coverage of confidence intervals for density. Overdispersion in the detection process may also reduce the coverage of confidence intervals, and we briefly consider how this might arise. One possibility is that stochasticity in the intensity surface may lead not only to overdispersion of *n*, but also to bias in the estimated variance of the detection parameters, and hence var[*a*(θ)] in Equation (1). We conducted simulations to test for an effect of stochastic AC generating processes on the coverage of confidence intervals for *a*(θ^^^), but found none (Appendix S5).

More interesting is the reverse possibility that non-independence in the detection process generates or amplifies overdispersion in *n*. We found evidence for a small effect of cohesion (synchronous detection of group members) on *n* (Appendix S5).

## Discussion

The Poisson point process model for activity centres in SECR may be strongly inhomogeneous without causing significant bias in estimates of sampling variance from homogeneous models (Fig. 2). This is expected because the number of detected individuals *n* remains Poisson under repeated sampling from a static inhomogeneous intensity surface i.e. *n* is not overdispersed.

However, stochastic variation in the intensity surface (vis. a Cox process) invariably results in overdispersion of *n* (Fig. 3) and poor coverage of confidence intervals with respect to the global density (Fig. 4, Table 1). The effect is less extreme for models fitted assuming a fixed population size in some area *A* around the detectors (Fig. 5), but this arbitrary assumption cannot be justified for common study designs as we discuss further below.

Whether users of SECR should be concerned about spatial overdispersion depends on their understanding of the population and the scope of inference. It will often be satisfactory to limit inference to the surveyed population, implicitly conditioning on the prevailing IHPP intensity surface, and then no overdispersion arises. Furthermore, a varying intensity surface that is fully determined by known covariates is not stochastic when these are included in the model (e.g., Diggle, 2003 p. 70), and no overdispersion is expected.

Problems arise when inference is required beyond the spatial or temporal scope of the sampling. We identify two such scenarios: clustering of AC in social species (Bischof et al. 2020) and extrapolation from spatial subsamples to a larger region.

### Clustered activity centres

Clustering may be treated mathematically as a Cox process, in which the intensity surface results from one realised set of ‘parent’ locations. To condition on such a surface seems not to make biological sense when the number, location and membership of clusters are dynamic. The scope of inference will then usually encompass different configurations. The sampling variance of global density is underestimated for populations with dynamic clusters, as recognised by Bischof et al. (2020). Our simulation results extend those of Bischof et al. (2020) to scenarios with random cluster size and non-zero within-cluster dispersion.

The variance inflation factor *c*^ of Bischof et al. (2020) uses internal evidence from one realisation of a Cox process to predict overdispersion of *n* across many realisations. This has proved to be inadequate, both empirically and from the simple logic that the generating process cannot be inferred from a static pattern. When AC come from an IHPP with static intensity rather than a Cox process, variance adjustment using *c*^ results in greater than nominal coverage and is therefore counterproductive.

We defined the local density *D*_*a*_ as a scalar-valued function, an integral of the IHPP intensity surface weighted by location-specific detection probability. For a Cox process *D*_*a*_ is a random variable. We see two uses for *D*_*a*_. The first is a general insight: when the intensity surface is stochastic, interval coverage for *D*^^^ remains good in simulations when each density estimate is compared to *D*_*a*_ for the intensity surface specific to the realisation. This is because *n* is not overdispersed for the IHPP in each realisation. Secondly, the value of var(*D*_*a*_) can be determined by simulation given a model for the Cox process and values for the detection parameters.

Externally acquired knowledge of the generating process is thereby transformed into a usable estimate of *c*_*n*_. This is plausible in the case of clustering, for which cluster size μ and within-cluster dispersion have biological interpretations, although field estimation will be difficult. Software is described in Appendix S6. Spatial replication could also be used to estimate the overdispersion in *n* due to clustering, but we have not seen examples.

We followed Bischof et al. (2020) in inflating the entire sampling variance of *D*^^^ by *c*^. It is more logical to multiply only the first variance component in Equation (1) by factor *c*_*n*_, which reduces the effect on var(*D*^^^). Overdispersion of AC slightly increased uncertainty regarding the detection process (var[*a*(θ^^^)]), but coverage of confidence intervals for *a*(θ^^^) was not impaired. If a suitable variance inflation factor is found for overdispersion due to non-independence and heterogeneity in the detection process then it should logically be applied to the second factor in Equation (1).

Complete cohesion of detection within spatial clusters is a special case in which the variance components are affected equally and inflation of var(*D*^^^) with the empirical measure *c*^ is justified. However, complete within-group cohesion seems unlikely if group members are dispersed in space. Cohesion causes overdispersion in detection itself (e.g., Appendix S5), which will be quantitatively important in studies where var[*a*(θ^^^)] is large.

### Spatial replication

It can be efficient to infer population density in a large region from a collection of small independent detector arrays (subgrids). The sampling variance of regional density then must allow for variation due to the placement of the subgrids relative to unknown variation in density within the region. We suggest viewing the regional population as a Cox process and applying a variance inflation factor for *D*^^^ that reflects variation in *D*_*a*_, the local density of the collection as a whole.

The sample variance of the number of individuals detected at each subgrid, divided by its mean, is a straightforward and effective estimate of overdispersion when subgrids are identical, independent and sufficiently numerous. Extension to other scenarios (e.g., subgrids of varying size or varying effort or that overlap in the individuals they sample) requires further work along the lines of Fewster et al. (2009), noting that they applied a different frame of reference (stochastic sampling locations instead of a stochastic point process for object locations).

Howe et al. (2022) performed simulations that appeared to refute the need for a variance adjustment. In their scenarios, three zones of differing density were sampled with two subgrids in each; coverage of unadjusted confidence intervals for global density was close to the nominal 95% when detection parameters did not vary. The simulations enforced a fixed density for each subgrid and the aggregate *D*_*a*_ for all subgrids was constant. Hence, as expected for a static IHPP, there was no overdispersion in *n*. When local densities were drawn at random from a distribution, as in a stochastic IHPP, the counts were overdispersed relative to a Poisson distribution and coverage was poor (Table 1). The adjustment for overdispersion was effective in terms of CI coverage, particularly when the number of subgrids was increased; precision was greatly reduced owing to the high variability of the simulated Cox process.

The general expression for overdispersion (Equation 4, replacing *n*_*k*_ with *n* _*j*_) allows the expected number on each subgrid E(*n*_*j*_) to vary according to a spatial model for density. We therefore expect the method to be useful when the fitted model includes habitat covariates and additional coefficients are estimated (*ν* > 1). The variance inflation factor *c*^_*n*_ would then apply to habitat-specific densities and to the regional population size *N*^^^ (e.g., Efford & Fewster, 2013).

### Fixed population size

It is common practice to compute the variance of *D*^^^ conditional on fixed *N* (*A*). This is implied in Bayesian estimation using data augmentation (Royle et al., 2014). Conditioning on *N* (*A*) is appealing because it shrinks the computed confidence intervals. For a Cox process, conditioning also severely constrains var(*D*_*a*_) and overdispersion is reduced, even relative to the lower (binomial) variance of *n* that applies when *N* (*A*) is fixed. The simulated coverage of variance-inflated intervals was improved, especially with data generated from a conditional Thomas process (Fig. 5).

However, conditioning on fixed *N* (*A*) is inelegant if the choice of *A* is arbitrary, and suggests an improbable scope of inference. It is not clear why we would want to restrict inference to the subset of realisations that result in one particular value of *N* (*A*). Specifically, it is biologically implausible that *N* (*A*) is fixed in IHPP scenarios with stochastically varying intensity or dynamic clustering. *N* (*A*) has been considered fixed when evaluating design-based (i.e. model-free) estimators of the encounter rate in distance sampling, reasoning that a fixed population of individuals might relocate themselves within *A* between realisations (Fewster et al., 2009). It is hard to reconcile this with biological models for the distribution of AC; as stated by Diana et al. (2022), “assuming that the number of points is independent of the spatial structure of the process is often unrealistic”.

## Data availability

Simulation results are archived in Zenodo at https://doi.org/10.5281/zenodo.15461025 (Efford 2025d). Efford (2025d) also includes an R script to generate the text figures and a vignette detailing the simulations (summaries for each scenario are in an appendix to the vignette). Other code required to perform the simulations is publicly available as the R package ‘overdispsim’ at https://github.com/MurrayEfford/overdispsim, archived in Zenodo at https://doi.org/10.5281/zenodo.15460402.

## Acknowledgments

We thank Adrian Baddeley and Tilman Davies for their help with the simulation of point processes. We thank two anonymous reviewers, Cyril Milleret and Eric Howe for their helpful comments on a draft.

## Funding

The authors received no funding for this work.

## Conflict of interest disclosure

The authors of this preprint declare that they have no conflict of interest relating to the content of this article.

## Supplementary information

The supplements archived with this preprint comprise the appendices:

Appendix S1. Examples of activity centres simulated with various Cox processes

Appendix S2. Derivation of expression for overdispersion leading to Equation 3

Appendix S3. Simulations of *c*_*n*_ and *c*^

Appendix S4. Tabular summary of simulations

Appendix S5. Overdispersion and the detection process

Appendix S6. Computation of overdispersion in *n* by simulation of random fields

## Notes

https://doi.org/10.5281/zenodo.15460402

https://doi.org/10.5281/zenodo.15460421

https://doi.org/10.5281/zenodo.15460409

### Competing Interest Statement

The authors have declared no competing interest.

### Summary of Updates

A badge has been added indicating that the preprint has been peer reviewed and recommended by PCI-Ecology Line numbers have been removed, line spacing reduced to 1.5, and a handful of typos and errors in reference formatting have been corrected.

