## Appendices for "Effect of spatial overdispersion on confidence intervals for population density estimated by spatial capture–recapture"

**Efford, M. G. and D. Fletcher. The effect of spatial overdispersion on confidence intervals for population density estimated by spatially explicit capture–recapture. BioRxiv**

### Appendix S1. Examples of activity centres simulated with various Cox processes

Three Cox processes were described in the main text: the Thomas cluster process, random-habitat IHPP, and log-Gaussian Cox processes. Figures on the following pages extend the examples in Fig. 1 of the main text to illustrate the effect of varying parameters. Rows represent levels of the parameter controlling inhomogeneity. Columns represent independent realisations (both the intensity surface and the point pattern simulated from that surface are random).

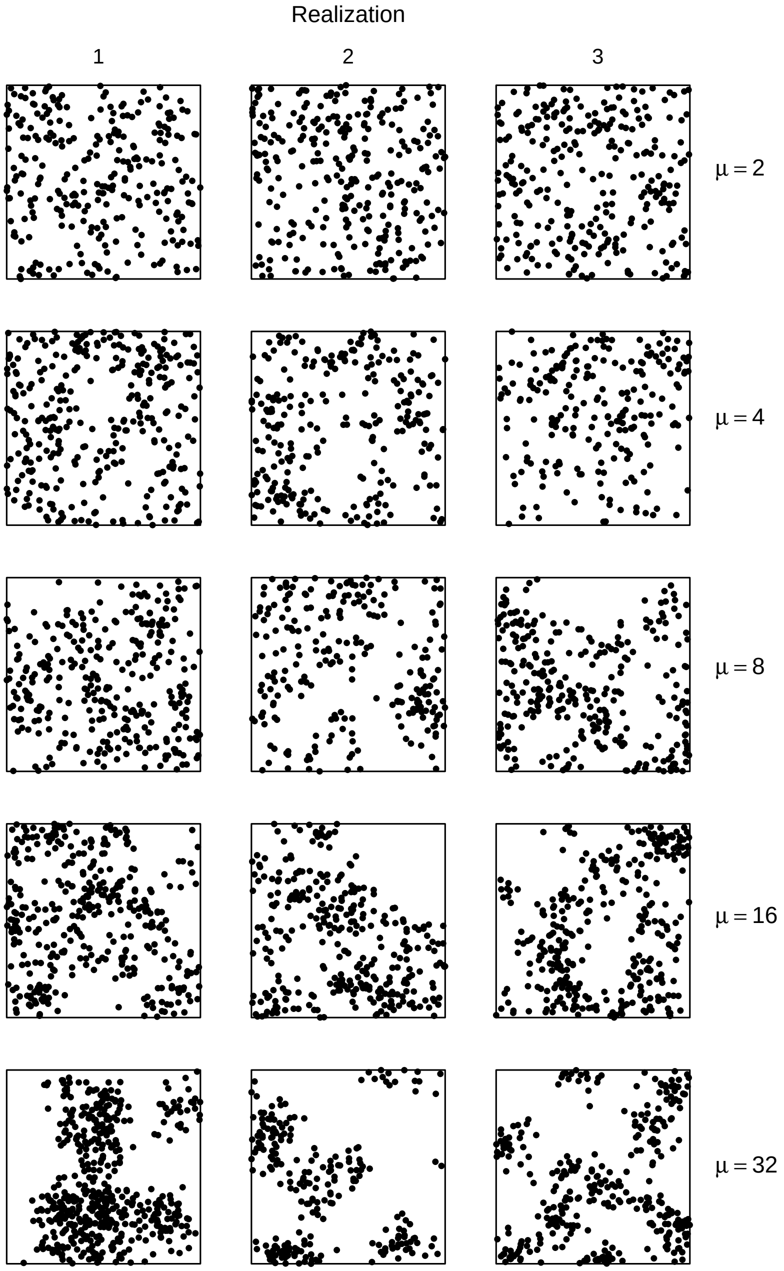


**Figure S1:** Simulated AC from Thomas (Neyman-Scott) cluster model in area $A$, a square $30\sigma \times 30\sigma$ ($\sigma$ defined in main text). Poisson-distributed parent points each give rise to a Poisson number of offspring (expectation $\mu \in \{2, 4, 8, 16, 32\}$) that follow circular bivariate normal distribution about the parent (scale $2\sigma$ in these examples). $\mathrm{E}[N(A)] = 256$.

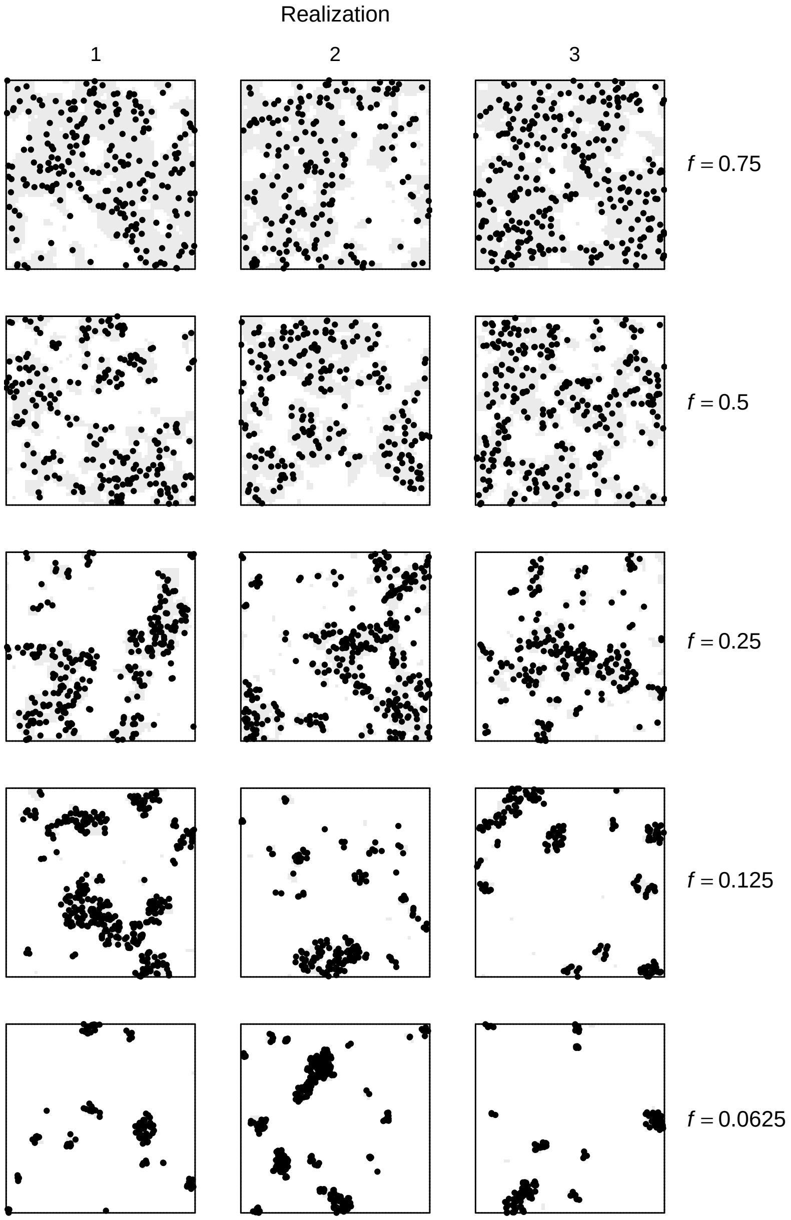


**Figure S2:** Simulated AC from random habitat IHPP model in area $A$, a square $30\sigma \times 30\sigma$ ($\sigma$ defined in main text). Landscape is a mosaic of habitat patches (grey shading) generated by method of Saura and Martínez-Millán (2000; fragmentation parameter $p = 0.5$). Habitat on average occupies fraction $f$ of frame. $\mathrm{E}[N(A)] = 256$.

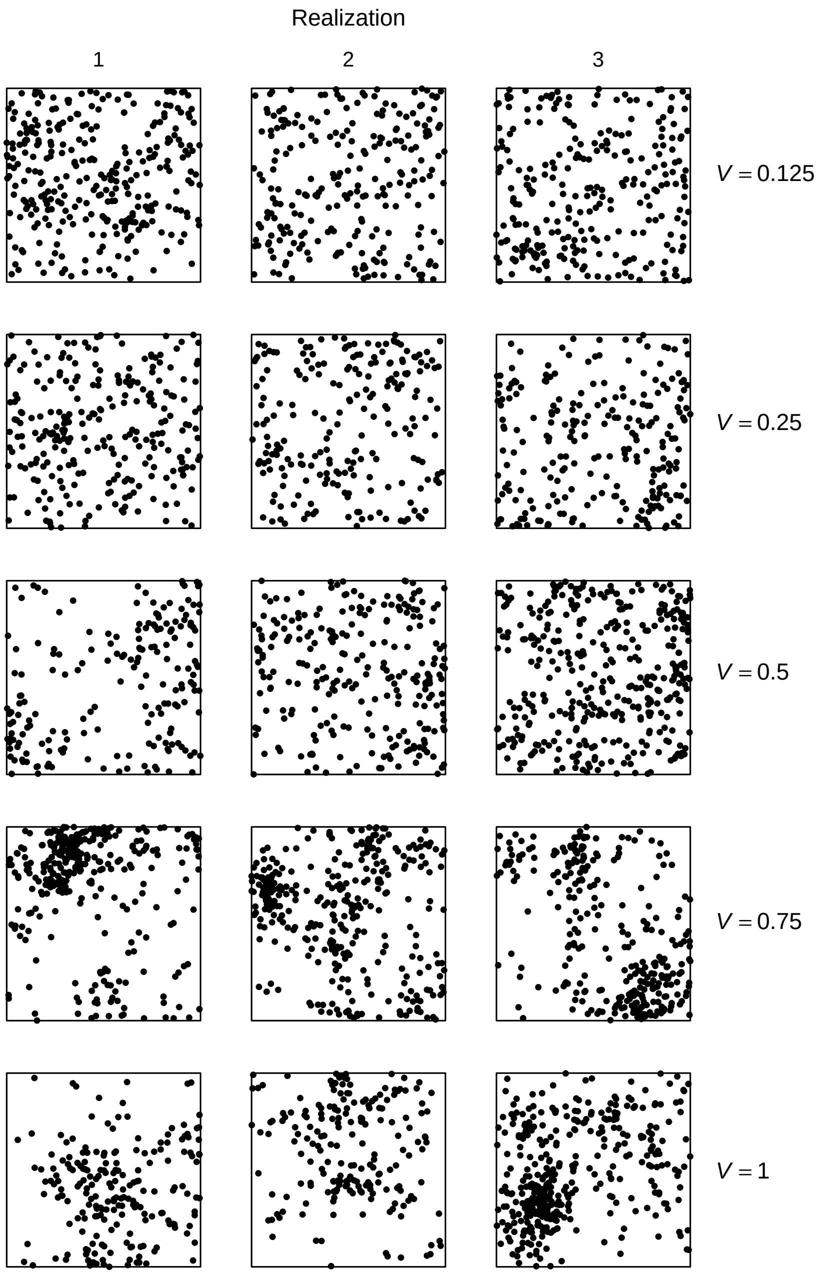


**Figure S3:** Simulated AC from LGCP model in area $A$, a square $30\sigma \times 30\sigma$ ($\sigma$ defined in main text). Gaussian random field with variance $V$ on log scale, and mean $\mu$ adjusted to keep $\mathrm{E}[N(A)] = 256$; exponential spatial covariance scale = $5\sigma$.

**Efford, M. G. and D. Fletcher. The effect of spatial overdispersion on confidence intervals for population density estimated by spatially explicit capture–recapture. BioRxiv**

### Appendix S2. Derivation of expression for overdispersion (Equation 3)

We can obtain an expression for the variance of $n$ in terms of both the variation in $D_a$ and the variation in $n$ for fixed $D_a$, as follows (Casella & Berger, 2002, Theorem 4.4.7):

$$\text{var}(n) = \text{E}_{D_a}[\text{var}_{n|D_a}(n|D_a)] + \text{var}_{D_a}[\text{E}_{n|D_a}(n|D_a)],$$

where a subscript indicates the distribution over which an expectation or variance is calculated.

As $n|D_a \sim$ Poisson, its mean and variance are equal and we have

$$\begin{aligned} \text{var}(n) &= \text{E}_{D_a}[\text{E}_{n|D_a}(n|D_a)] + \text{var}_{D_a}[\text{E}_{n|D_a}(n|D_a)] \\ &= \text{E}(n) + \text{var}_{D_a}[\text{E}_{n|D_a}(n|D_a)], \end{aligned}$$

since $\text{E}(n) = \text{E}_{D_a}[\text{E}_{n|D_a}(n|D_a)]$ (Casella & Berger, 2002, Theorem 4.4.3).

By definition of $D_a$, we have

$$\text{E}_{n|D_a}(n|D_a) \equiv \int_A D(\mathbf{x}) p_\cdot(\mathbf{x};\theta) d\mathbf{x} = aD_a.$$

This leads to

$$\text{var}(n) = \text{E}(n) + a(\theta)^2 \,\text{var}_{D_a}(D_a),$$

with $\text{E}(n) = a(\theta)\text{E}_{D_a}(D_a)$, as in Equation (3) (in which the subscript $D_a$ is omitted for simplicity).

**Efford, M. G. and D. Fletcher. The effect of spatial overdispersion on confidence intervals for population density estimated by spatially explicit capture–recapture. BioRxiv**

### Appendix S3. Simulations of $c_n$ and $\hat{c}$

The simulations described here were used to evaluate $\hat{c}$ (the variance inflation factor of Bischof et al. (2020)) as an estimate of the empirical overdispersion of $n$. These simulations did not require any fitting the SECR model.

The three Cox processes for generating distributions of activity centers (AC) were described in the main text. Additional scenarios were simulated (Table S1). All simulations used an area 30 units square with an expected population of 256 within the area (0.2844444/unit$^2$).

**Table S1:** Simulation parameters

| Cox process | Parameter values | Description |
|---|---|---|
| Thomas cluster | $\mu$ = 1, 2, 4, 8, 16, 32 | expected number per parent |
| | scale = 0.0001, 1, 2, 4 | dispersion around parent |
| Random habitat | f = 0.0625, 0.125, 0.25, 0.5, 0.75, 1 | habitat fraction |
| | p = 0.25, 0.5 | fragmentation parameter |
| LGCP | $V$ = 0, 0.125, 0.25, 0.5, 0.75, 1 | variance of random field |
| | scale = 2, 5, 10 | scale of exponential covariance |

Results of these simulations are plotted in Fig. S1.

We also calculated $c_n = 1 + a(\theta)\ \mathrm{var}(D_a)/\mathrm{mean}(D_a)$ from replicate simulations of the intensity surface to validate $c_n$ as a substitute for full simulations of the empirical overdispersion of $n$ (i.e. without simulating AC realisations and the detection process). See main text for notation and $\theta$. The resulting $c_n$ were almost exactly equal to the empirical overdispersion of $n$ (Fig. S2).

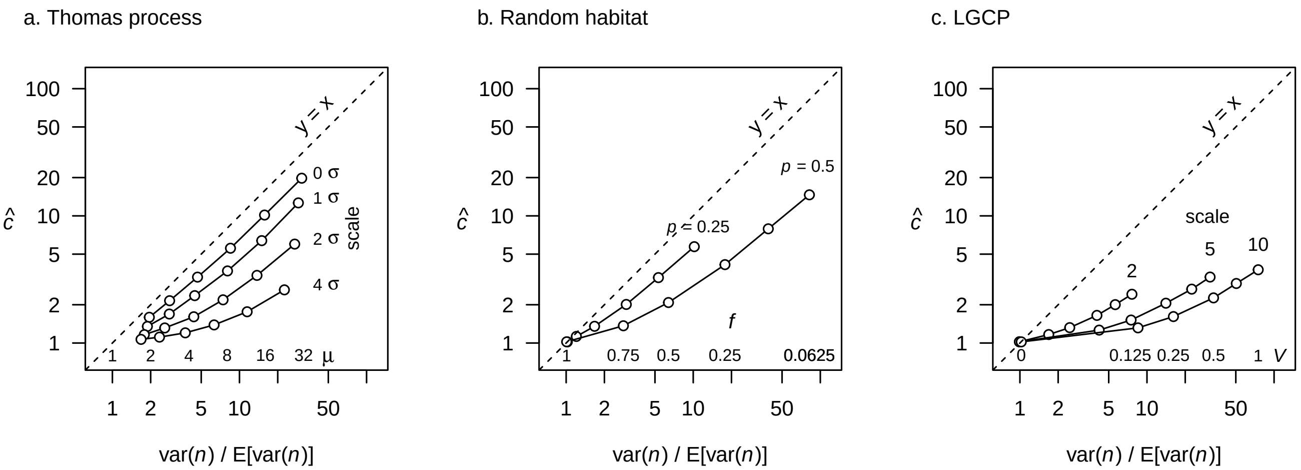


**Figure S1:** Effect of stochastic AC generating processes on the relationship between the overdispersion of $n$ and an empirical measure of overdispersion $\hat{c}$. (a) Thomas (Neyman-Scott) distributions with various parameter levels, (b) random habitat, and (c) Log-Gaussian Cox process with variance $V$ and three levels of scale. Note log scale on both axes. 10000 simulations per scenario

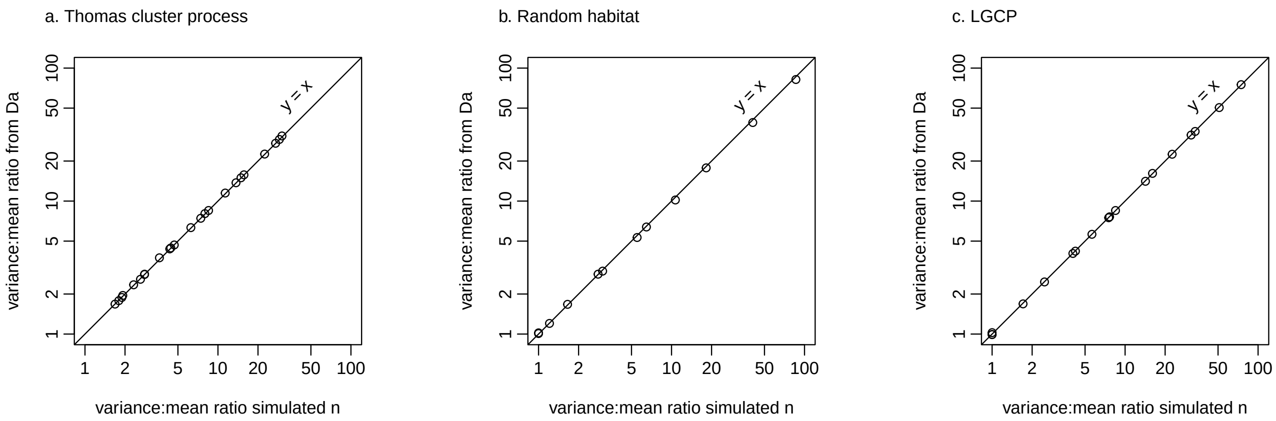


**Figure S2:** Validation of $c_n$ by comparison with simulated overdispersion of $n$. Each point represents a scenario from Table S1. 10000 simulations per scenario.

**Efford, M. G. and D. Fletcher. The effect of spatial overdispersion on confidence intervals for population density estimated by spatially explicit capture–recapture. BioRxiv**

### Appendix S4. Tabular summary of results

This Appendix gives numerical results for the simulations presented graphically in the main text. These assess estimates of average population density $\hat{D}$ from fitting a homogeneous-density model to data generated by three Cox processes, i.e. stochastic-intensity inhomogeneous Poisson point processes. The 'true' density used to assess relative bias and coverage of confidence intervals could be either the global mean of the Cox process (Table S1) or the local density in the vicinity of detectors, as defined in the main text (Table S2).

The fields in the following tables are

| | |
|---|---|
| $f$ | fraction of landscape in habitat patches |
| $V$ | variance of Gaussian field in LGCP generating model for AC |
| $\mu$ | expected number of AC per cluster for Thomas process |
| RB | relative bias of density estimates |
| COV | coverage of naive 95% confidence intervals |
| $\hat{c}$ | overdispersion computed from number of individuals at each detector $k$ |
| $\mathrm{var}(n)/\mathrm{var}_0(n)$ | variance of $n$ relative to fitted model |
| Adjusted COV | coverage of confidence intervals using $\hat{c}$ as variance inflation factor |

**Table S1:** Simulation results relative to global density

Thomas cluster process

| | $\mu$ | | | | | |
|---|---|---|---|---|---|---|
| | 1 | 2 | 4 | 8 | 16 | 32 |
| RB | -0.001 | 0.000 | 0.000 | 0.010 | -0.010 | -0.005 |
| COV | 0.857 | 0.783 | 0.680 | 0.508 | 0.394 | 0.293 |
| $\hat{c}$ | 1.128 | 1.282 | 1.524 | 2.042 | 3.148 | 5.224 |
| $\text{var}(n)/\text{var}_0(n)$ | 1.747 | 2.607 | 4.133 | 7.391 | 13.950 | 27.002 |
| Adjusted COV | 0.889 | 0.838 | 0.758 | 0.688 | 0.636 | 0.597 |

Random habitat

| | $f$ | | | | | |
|---|---|---|---|---|---|---|
| | 0.0625 | 0.125 | 0.250 | 0.500 | 0.750 | 1.000 |
| RB | 0.0390 | 0.021 | 0.019 | 0.004 | -0.007 | 0.001 |
| COV | 0.1600 | 0.230 | 0.354 | 0.545 | 0.730 | 0.945 |
| $\hat{c}$ | 12.5560 | 7.110 | 3.783 | 1.955 | 1.333 | 1.008 |
| $\text{var}(n)/\text{var}_0(n)$ | 79.9850 | 41.231 | 17.361 | 6.773 | 3.023 | 1.049 |
| Adjusted COV | 0.5610 | 0.601 | 0.629 | 0.710 | 0.795 | 0.950 |

LGCP

| | $V$ | | | | | |
|---|---|---|---|---|---|---|
| | 0.000 | 0.125 | 0.250 | 0.500 | 0.750 | 1.000 |
| RB | -0.002 | 0.003 | -0.014 | 0.002 | 0.030 | 0.009 |
| COV | 0.955 | 0.514 | 0.374 | 0.314 | 0.244 | 0.189 |
| $\hat{c}$ | 1.008 | 1.170 | 1.376 | 1.718 | 2.095 | 2.558 |
| $\text{var}(n)/\text{var}_0(n)$ | 0.962 | 8.499 | 15.007 | 27.571 | 47.199 | 72.479 |
| Adjusted COV | 0.958 | 0.544 | 0.430 | 0.396 | 0.327 | 0.302 |

**Table S2:** Simulation results relative to detection-weighted local density

Thomas cluster process

| | $\mu$ | | | | | |
|---|---|---|---|---|---|---|
| | 1 | 2 | 4 | 8 | 16 | 32 |
| RB | 0.000 | 0.001 | 0.000 | -0.004 | 0.000 | -0.002 |
| COV | 0.952 | 0.938 | 0.954 | 0.951 | 0.954 | 0.960 |
| $\hat{c}$ | 1.144 | 1.267 | 1.552 | 2.062 | 3.119 | 5.132 |
| $\text{var}(n)/\text{var}_0(n)$ | 1.796 | 2.620 | 4.285 | 7.665 | 14.851 | 27.415 |
| Adjusted COV | 0.966 | 0.964 | 0.986 | 0.995 | 0.998 | 0.999 |

Random habitat

| | $f$ | | | | | |
|---|---|---|---|---|---|---|
| | 0.0625 | 0.125 | 0.250 | 0.500 | 0.750 | 1.000 |
| RB | -0.0030 | -0.001 | -0.002 | -0.001 | 0.000 | 0.001 |
| COV | 0.9500 | 0.943 | 0.944 | 0.943 | 0.945 | 0.945 |
| $\hat{c}$ | 12.5560 | 7.110 | 3.783 | 1.955 | 1.333 | 1.008 |
| $\text{var}(n)/\text{var}_0(n)$ | 79.9850 | 41.231 | 17.361 | 6.773 | 3.023 | 1.049 |
| Adjusted COV | 1.0000 | 1.000 | 0.999 | 0.988 | 0.967 | 0.950 |

LGCP

| | $V$ | | | | | |
|---|---|---|---|---|---|---|
| | 0.000 | 0.125 | 0.250 | 0.500 | 0.750 | 1.000 |
| RB | -0.002 | 0.001 | -0.004 | 0.003 | 0.001 | 0.001 |
| COV | 0.955 | 0.940 | 0.943 | 0.953 | 0.943 | 0.951 |
| $\hat{c}$ | 1.008 | 1.170 | 1.376 | 1.718 | 2.095 | 2.558 |
| $\text{var}(n)/\text{var}_0(n)$ | 0.962 | 8.499 | 15.007 | 27.571 | 47.199 | 72.479 |
| Adjusted COV | 0.958 | 0.956 | 0.975 | 0.981 | 0.981 | 0.990 |

### Appendix S5. Overdispersion and the detection process

The main text focuses on the state model (distribution of activity centers AC) and overdispersion of the number of individuals detected $n$ relative to the expected variance of a Poisson or binomial distribution. The detection process also contributes to the sampling variance of $\hat{D}$ because of uncertainty in the effective sampling area $a(\hat{\theta})$ (main text Equation (1)). Here we consider the impact of spatial overdispersion on $a(\hat{\theta})$, and the effect of overdispersion in the detection process itself on $n$ and $a(\hat{\theta})$.

#### Spatial overdispersion and the effective sampling area $a(\hat{\theta})$

The estimated effective sampling area $a(\hat{\theta})$ is a scalar value that encapsulates the detection process in spatial capture–recapture models, closely analogous to detection probability in non-spatial capture–recapture models. Unbiased estimation of $a$ and its sampling variance is required for estimation of average density. We considered whether spatial overdispersion (a Cox process for AC) might affect estimates of detection parameters so as to inflate $\widehat{\text{var}}[a(\hat{\theta})]$ in Equation (1) of the main text.

We used $\widehat{\text{var}}[a(\hat{\theta})] = \hat{G}_\theta \hat{I}_\theta \hat{G}_\theta^T$, where $\hat{I}_\theta$ is the information matrix (inverse Hessian) of the detection parameters evaluated at the maximum likelihood estimates, and $\hat{G}_\theta$ is a numerical estimate of the gradient of $a(\theta)$ with respect to each of the detection parameters. We evaluated the coverage of a 95% symmetric Wald interval using this sampling variance. Simulation conditions followed the main text. Code is provided in Efford (2025a).

We found no evidence that spatial overdispersion itself inflated the variance of $a(\hat{\theta})$: coverage of confidence intervals for $a(\hat{\theta})$ was close to the nominal level (Table S1).

**Table S1:** Simulation results: effective sampling area $a$

Thomas cluster process

| | $\mu$ | | | | | |
|---|---|---|---|---|---|---|
| | 1 | 2 | 4 | 8 | 16 | 32 |
| RB | 0.001 | 0.000 | 0.000 | 0.001 | 0.001 | 0.000 |
| RSE | 0.009 | 0.009 | 0.009 | 0.009 | 0.009 | 0.010 |
| COV | 0.950 | 0.951 | 0.947 | 0.943 | 0.958 | 0.948 |

Random habitat

| | $f$ | | | | | |
|---|---|---|---|---|---|---|
| | 0.0625 | 0.125 | 0.250 | 0.500 | 0.750 | 1.000 |
| RB | 0.000 | 0.000 | 0.000 | 0.000 | 0.000 | 0.000 |
| RSE | 0.012 | 0.010 | 0.009 | 0.009 | 0.009 | 0.009 |
| COV | 0.919 | 0.922 | 0.938 | 0.954 | 0.944 | 0.948 |

LGCP

| | $V$ | | | | | |
|---|---|---|---|---|---|---|
| | 0.000 | 0.125 | 0.250 | 0.500 | 0.750 | 1.000 |
| RB | 0.000 | 0.001 | 0.000 | 0.000 | 0.000 | 0.000 |
| RSE | 0.009 | 0.009 | 0.009 | 0.009 | 0.010 | 0.010 |
| COV | 0.941 | 0.939 | 0.953 | 0.953 | 0.938 | 0.928 |

Thomas cluster process with complete cohesion

| | $\mu$ | | | | | |
|---|---|---|---|---|---|---|
| | 1 | 2 | 4 | 8 | 16 | 32 |
| RB | 0.002 | 0.001 | 0.003 | 0.001 | 0.005 | 0.017 |
| RSE | 0.009 | 0.009 | 0.009 | 0.009 | 0.010 | 0.012 |
| COV | 0.831 | 0.759 | 0.646 | 0.497 | 0.363 | 0.271 |

#### Cohesion

One source of overdispersion in the detection process was identified by Bischof et al. (2020) as 'cohesion' - a tendency of animals belonging to a particular group to be detected together. In simulations of a fixed cluster process they varied a cohesion parameter $\gamma$ between zero (independent detection) and one (complete synchrony of detection). We reproduced their results with function sim.cohesion () in the R package 'overdispsim' (Efford, 2025b), used in conjunction with the R package 'secrdesign' (Efford, 2025c). For simplicity we consider only complete cohesion ($\gamma = 1$); results are intermediate for intermediate levels of $\gamma$. The original definition of cohesion assumed negligible dispersion of AC within clusters (zero scale), and would appear less meaningful or realistic when AC are dispersed from the parent location. We therefore consider only scenarios with negligible scale (0.0001 for the rThomas function of spatstat that does not allow scale = 0).

##### Effect of cohesion on $n$

Cohesion increases overdispersion in $n$ and reduces coverage of confidence intervals for $\hat{D}$, as reported by Bischof et al. (2020). However, we found the effect on overdispersion of $n$ to be quite small for the scenarios we considered (Fig. S1). For the fixed-cluster-size process (Fig. S1b) with complete cohesion, parents are the only independent units: the dataset is obtained by cloning a detection history simulated for each notional parent a multiple of $\mu$ times, where $\mu$ is the cluster size, and the resulting overdispersion of $n$ is exactly $\mu$ (cf Anderson et al., 1994). With clusters of fixed size and complete cohesion the empirical estimate $\hat{c}$ was an unbiased estimate of cluster size and therefore of $c_n$ (Fig. S2).

##### Effect of cohesion on $a(\hat{\theta})$

Further simulations with a Thomas process for AC and complete within-cluster cohesion of detection (Table S1) show that the principal effect of cohesion is on estimates of the detection parameters themselves, as might be expected. We expect this to manifest as underestimation of the sampling variance of $\hat{D}$, in proportion to the contribution of $a(\hat{\theta})$. In the scenario used in our simulations the contribution was small ($\approx 1\%$) so we do not pursue the matter. In real world scenarios the effect may be dominant.

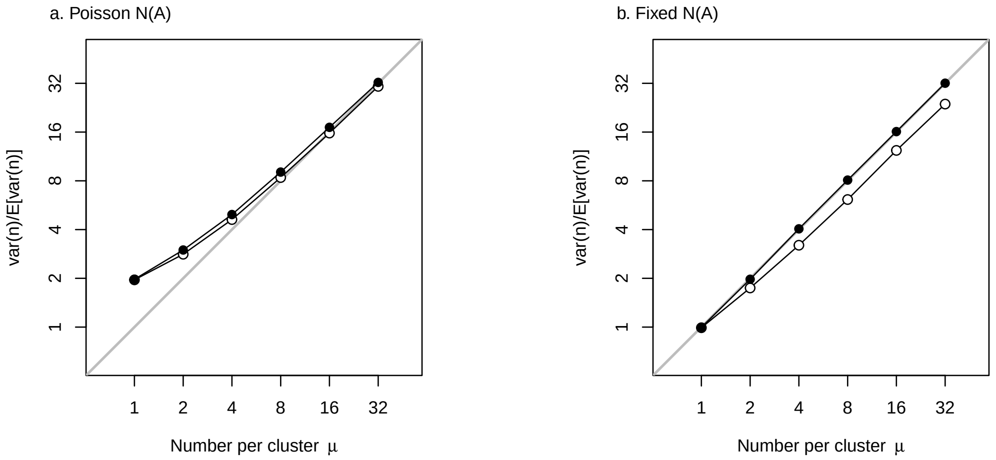


**Figure S1:** Overdispersion of $n$ from cluster model in relation to cluster size $\mu$ and cohesion. Cohesion was either zero (open dots) or complete (filled dots). (a) Poisson number of parents and Poisson cluster size (Thomas distribution) (b) fixed number of parents, fixed cluster size.

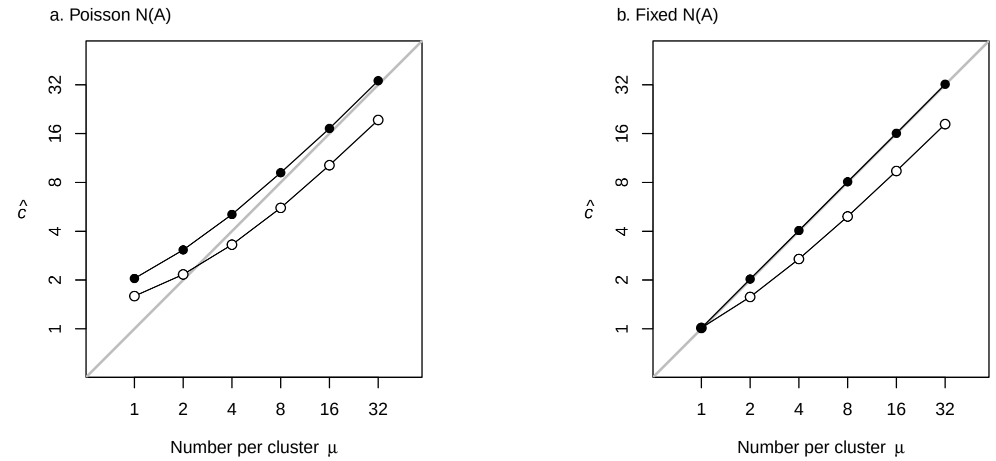


**Figure S2:** Estimated overdispersion $\hat{c}$ of $n$ from cluster model in relation to cluster size $\mu$ and cohesion. Cohesion was either zero (open dots) or complete (filled dots). (a) Poisson number of parents and Poisson cluster size (Thomas distribution) (b) fixed number of parents, fixed cluster size.

### Appendix S6. Computation of overdispersion in *n* by simulation of random fields

Overdispersion in $n$ for a particular Cox process with known parameters may be calculated directly from realisations of the intensity surface as described in the main text. Code for simulation-based estimation of $c_n$ is provided in the R package **secrRFS** available on GitHub and archived on Zenodo (Efford, 2025).

A simple application of **secrRFS** is demonstrated below.

```
library(secrRFS)
detectpar <- list(lambda0 = 0.5, sigma = 1)
grid144 <- make.grid(12,12, detector = 'proximity', spacing = 2.0)
grid144mask <- make.mask(grid144, spacing = 0.5, buffer = 4)
D <- 256/maskarea(grid144mask)
parm <- list(D = D, mu = 8, scale = 2)
RFS(randomfn = randomParents,  parm, nrepl = 1000,  traps = grid144, mask =
    grid144mask, detectfn = 'HHN', detectpar = detectpar, noccasions = 5)
[1] 7.638902
```

The result is a value for $c_n$ that may be used as a variance inflation factor, assuming in this case a Thomas process with 8 expected offspring per parent and scale 2.

Another example follows the stochastic version of Howe et al. (2022) Scenario 2 (Table 1 in main text). We first define a function to select parm$ngrids densities at random with replacement from vector of values parm$D:

```
Dfun <- function (mask, parm, plt = FALSE) {
    D <- sample (parm$D, size = parm$ngrids, replace = TRUE)
    rep(sum(D), nrow(mask))
}
trps       <- make.grid(5,8, detector = 'proximity', spacing = 2000)
msk        <- make.mask(trps, buffer = 8000)
```

```
RFS(Dfun, parm = list(D = c(6,18,12)*1e-4, ngrids = 6), nrepl = 100000,
        traps = trps, mask = msk, detectfn = 'HN', detectpar = list(g0 = 0.3,
        sigma = 1500), noccasions = 6, verbose = FALSE)
[1] 6.034851
```

The resulting $c_n$ is close to the mean of many empirical estimates in Table 1 of the main text.

The function `carray` in **secrRFS** may also be used to generate tables such as these:

```
parmlevels <- list(D = D, mu = 2^(0:5), scale = c(1e-04, 1, 2, 4, 8))
RFSargs <- list (randomfn = randomParents, nrepl = 1000, traps  = grid144, mask =
    grid144mask,  detectfn   = 'HHN',  detectpar  = detectpar, noccasions = 5)
ca <- carray (parmlevels, RFSargs)
round(ca, 1)

    scale
mu   1e-04    1    2    4    8
  1    1.9  1.8  1.8  1.7  1.4
  2    2.8  2.8  2.6  2.3  1.8
  4    4.8  4.6  4.1  3.6  2.5
  8    8.3  8.3  7.4  6.5  4.3
  16  16.5 15.3 13.4 11.9  7.4
  32  30.3 28.6 26.0 22.3 14.2
```

The entries in the table are the required variance inflation factors when the distribution of AC results from a Thomas cluster process with parameters ‘mu’ and ‘scale’.
